# Diaphorin, a polyketide produced by a bacterial symbiont of the Asian citrus psyllid, inhibits the growth of *Bacillus subtilis* but promotes the growth of *Escherichia coli*

**DOI:** 10.1101/2022.05.09.491270

**Authors:** Nozomu Tanabe, Rena Takasu, Yuu Hirose, Yasuhiro Kamei, Maki Kondo, Atsushi Nakabachi

## Abstract

Diaphorin is a polyketide produced by *Candidatus* Profftella armatura (Gammaproteobacteria: Burkholderiales), an obligate symbiont of a notorious agricultural pest, the Asian citrus psyllid *Diaphorina citri* (Hemiptera: Psyllidae). Diaphorin belongs to the pederin family of bioactive agents found in various host-symbiont systems, including beetles, lichens, and sponges, harboring phylogenetically diverse bacterial producers. Previous studies showed that diaphorin has inhibitory effects on various eukaryotes, including the natural enemies of *D. citri*. However, little is known about its effects on prokaryotic organisms. To address this issue, the present study assessed the biological activities of diaphorin on two model prokaryotes, *Escherichia coli* (Gammaproteobacteria: Enterobacterales) and *Bacillus subtilis* (Firmicutes: Bacilli). The analyses revealed that diaphorin inhibits the growth of *B. subtilis* but moderately promotes the growth of *E. coli*. This finding implies that diaphorin functions as a defensive agent of the holobiont (host + symbionts) against some bacterial lineages but is beneficial for others, which potentially include obligate symbionts of *D. citri*.

**Importance:** Certain secondary metabolites, including antibiotics, evolve to mediate interactions among organisms. These molecules have distinct spectra for microorganisms and are often more effective against Gram-positive bacteria than Gram-negative ones. However, it is rare that a single molecule has completely opposite activities on distinct bacterial lineages. The present study revealed that a secondary metabolite synthesized by an organelle-like bacterial symbiont of psyllids inhibits the growth of Gram-positive *Bacillus subtilis* but promotes the growth of Gram-negative *Escherichia coli*. This finding not only provides insights into the evolution of symbiosis between animal hosts and bacteria but may also potentially be exploited to promote the effectiveness of industrial material production by microorganisms.

## Introduction

Microorganisms produce diverse secondary metabolites that mediate competition, communication, and other interactions with surrounding organisms and the environment (1–4). Such molecules have various biological activities, some of which facilitate symbiosis between microorganisms and animal hosts (5–8).

The Asian citrus psyllid *Diaphorina citri* Kuwayama (Hemiptera: Sternorrhyncha: Psylloidea: Psyllidae) is an important agricultural pest that transmits *Candidatus* Liberibacter spp. (Alphaproteobacteria: Rhizobiales), the causative agents of a devastating citrus disease known as huanglongbing (HLB) or greening disease (9– 12). Because HLB is currently incurable, controlling *D. citri* is presently the most crucial part of HLB management (9, 12). Although the application of chemical insecticides is currently the primary option for controlling *D. citri*, a more sustainable strategy, including biological control, is warranted (9, 13–15) partly due to the global increase in the resistance of *D. citri* to various pesticides (16–18).

The *D. citri* hemocoel contains a symbiotic organ called the bacteriome (19, 20), which harbors two distinct intracellular symbionts, *Ca*. Carsonella ruddii (Gammaproteobacteria: Oceanospirillales) and *Ca*. Profftella armatura (Gammaproteobacteria: Burkholderiales) (21, 22). *Carsonella* is a typical nutritional symbiont, providing its host with essential amino acids that are scarce in the phloem sap diet (21, 23, 24). In contrast, *Profftella* appears to be an organelle-like defensive symbiont, producing toxins that protect the holobiont (host + symbionts) from natural enemies (21, 25). *Profftella* has a very small genome at 460 kb, a large part of which is devoted to a gene set to synthesize a polyketide, diaphorin (21). Diaphorin is an analog of pederin (21), a defensive polyketide that accumulates in the body fluid of *Paederus* rove beetles (Coleoptera: Staphylinidae) to deter predators (26–28). Previous studies have demonstrated that diaphorin, which is contained at the concentration of 2 to 20 mM in *D. citri* depending on its life stage (29), has inhibitory effects on various eukaryotic organisms, suggesting that it helps protect *D. citri* from eukaryotic predators, parasitoids, parasites, and pathogens (21, 25, 30). Recent studies have revealed that *Profftella* and its gene clusters for synthesizing diaphorin are conserved in relatives of *D. citri*, suggesting the physiological and ecological importance of diaphorin for the host insect (31, 32). However, little is known about the effects of diaphorin on prokaryotic organisms.

To address this issue, this study assessed the biological activities of diaphorin on *Escherichia coli* (Gammaproteobacteria: Enterobacterales) and *Bacillus subtilis* (Firmicutes: Bacilli), which are model organisms for Gram-negative and Gram-positive bacteria, respectively.

## Materials and methods

### Preparation of diaphorin

Diaphorin was extracted and purified as described previously (21, 25). Adult *D. citri* were ground in methanol, and the extracts were concentrated *in vacuo*. The residue was purified in a Shimadzu (Kyoto, Japan) LC10 high-performance liquid chromatography (HPLC) system with an Inertsil ODS-3 C18 reverse-phase preparative column (GL Science, Tokyo, Japan). The purified samples were combined, dried, redissolved in methanol, and filter-sterilized using a Minisart syringe filter with a pore size of 0.2 μm (Sartorius, Göttingen, Germany). Aliquots of the purified samples were quantified in the LC10 HPLC system using an Inertsil ODS-3 analytical column (GL Science). The purified diaphorin was stored at −20°C until use.

### Transformation of *E. coli*

To confer ampicillin resistance and *β*-galactosidase activity, *E. coli* strain JM109 was transformed with the pGEM-T Easy Vector (Promega, Madison, WI, USA) that encodes *β*-lactamase and the *β*-galactosidase *α*-peptide (LacZα). After self-ligation with T4 DNA ligase at 25°C for 1 h, the vector was introduced into *E. coli* according to the manufacturer’s instructions. The nucleotide sequence of *lacZα* was checked following colony PCR using primers lacZ_F (5′-GCGCTGGCAAGTGTAGCGG-3′) and lacZ_R (5′-TCCGGCTCGTATGTTGTGTGG-3′), which respectively target the 5′ and 3′ flanking regions of the gene. Clones with intact *lacZα* lacking insertions due to T-overhangs were selected and used for the following assays.

### Evaluation of the effects of diaphorin on *E. coli*

*E. coli* cells transformed with the pGEM-T Easy plasmid were precultured in Luria-Bertani (LB) medium (1% Bacto tryptone, 0.5% Bacto yeast extract, and 1% NaCl, pH 7.0) containing 100 μg/mL ampicillin at 37°C for 14 h with reciprocal shaking (130 rpm). Growth was monitored by measuring the optical density of cultures at 600 nm (OD_600_) with a NanoDrop 2000c spectrophotometer (Thermo Fisher Scientific), with a 1 mm pathlength. Various diaphorin concentrations (5 μM–5 mM) were prepared in LB medium containing 100 μg/mL ampicillin. Precultured *E. coli* cells were inoculated into the diaphorin-containing medium, diluting the preculture at 1:1,000, and cultured for 24 h as before. The cell density of each culture was analyzed by measuring the OD_600_ as described above. Growth analyses were accompanied by controls cultured in the absence of diaphorin. Four temporally independent experiments were performed, each consisting of three independent cultures in three independent tubes per treatment, giving 12 independent cultures (n = 12) per treatment. To assess the direct effects of diaphorin on the optical densities of culture media, time-course analyses of OD_600_ of sterile (no inoculation of *E. coli*) medium containing 5 mM diaphorin were also performed at 37°C (n = 3).

### Transformation of *B. subtilis*

To confer tetracycline resistance, *B. subtilis* strain ISW1214 was transformed with the pHY300PLK (Takara, Kusatsu, Japan) plasmid that encodes a tetracycline resistance gene. Transformation of competent *B. subtilis* cells was performed using plasmids preamplified in *E. coli* strain BL21 (DE3) according to the manufacturer’s instructions.

### Evaluation of the effects of diaphorin on *B. subtilis*

*B. subtilis* cells transformed with the pHY300PLK plasmid were precultured in L-broth (1% Bacto tryptone, 0.5% Bacto yeast extract, and 0.05% NaCl, pH7.0) containing 20 μg/mL tetracycline at 37°C for 14 h with reciprocal shaking (130 rpm). Growth was monitored by measuring the OD_600_ as described above. Various diaphorin concentrations (5 μM–5 mM) were prepared in L-broth containing 20 μg/mL tetracycline. Precultured *B. subtilis* cells were inoculated to the diaphorin-containing medium, diluting the preculture at 1:1,000, and cultured for 24 h as before. The cell density of each culture was analyzed by measuring the OD_600_. Growth analyses were accompanied by controls cultured in the absence of diaphorin. Four temporally independent experiments were performed, each consisting of three independent cultures in three independent tubes per treatment, giving 12 independent cultures (n = 12) per treatment. To assess the direct effects of diaphorin on optical densities of culture media, time-course analyses of OD_600_ of sterile (no inoculation of *B. subtilis*) medium containing 5 mM diaphorin were also performed at 37°C (n = 3).

### Assessment of culture purity by amplicon sequencing

To assess the possibility of contamination, bacterial populations in culture media were analyzed using high-throughput amplicon sequencing of the 16S rRNA gene. After cultivating *E. coli* or *B. subtilis* with or without treatment of 5 mM diaphorin for 24 h, cells were harvested by centrifugation at 16,000 × *g* for 5 min. Cell pellets were resuspended in suspension buffer, which was transferred into NucleoSpin bead tubes type B containing 40 to 400 μm glass beads (Macherey-Nagel, Düren, Germany). The bead tubes were attached to Vortex-Genie 2 (Scientific Industries, Bohemia, NY, USA) using an MN bead tube holder, and cells were disrupted by agitation at 3,200 rpm for 20 min. Subsequently, DNA was extracted using NucleoSpin Microbial DNA columns according to the manufacturer’s instructions. Amplicon PCR was performed using extracted DNA, the KAPA HiFi HotStart ReadyMix (KAPA Biosystems, Wilmington, MA, USA), and the primer set 16S_341F (5′-TCGTCGGCAGCGTCAGATGTGTATAAGAGACAGCCTACGGGNGGCWGC AG-3′) and 16S_805R (5′-GTCTCGTGGGCTCGGAGATGTGTATAAGAGACAGGACTACHVGGGTATC TAATCC-3′) targeting the V3 and V4 regions of the 16S rRNA gene, based on the instructions by Illumina (San Diego, CA, USA) (33). Dual indices and Illumina sequencing adapters were attached to the amplicons with index PCR using the Nextera XT Index Kit v2 (Illumina). The libraries were combined with PhiX control version 3 (Illumina), and 300 bp of both ends were sequenced on the MiSeq platform (Illumina) with the MiSeq Reagent Kit version 3 (600 cycles; Illumina). After the amplicon sequence reads were demultiplexed, the output sequences were imported into the QIIME2 platform (version 2020.2) (34) and processed as described previously (31, 35). Obtained sequence variants were manually checked by performing BLASTN searches against the National Center for Biotechnology Information nonredundant database (36).

### Optical microscopic analysis

Aliquots of bacterial cultures were put on glass slides, stained with NucBlue Live ReadyProbes Reagent (Hoechst 33342 dye, Thermo Fisher Scientific) as needed, and examined by differential interference contrast (DIC) microscopy and/or fluorescence microscopy using the BX53 biological microscope (Olympus, Tokyo, Japan). The morphology of bacterial cells was analyzed using the Fiji package of ImageJ (37). The cell length (major axis) and cell width (minor axis) were measured using the segmented line tool implemented in ImageJ. In this study, even when septa or septa-like structures were observed, a sequential unit was defined as a single cell, if cleavage had not occurred. Cell volume was calculated assuming that cells consist of a cylinder and two half-spheres:

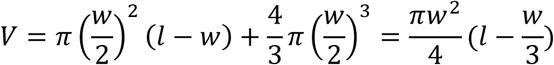

where l = cell length and w = cell width.

Aliquots of bacterial culture were put into a bacterial counter (depth of 20 μm; Sunlead Glass, Koshigaya, Japan), and cell numbers were counted under a BX53 microscope.

### Electron microscopic analysis

Cultured *E. coli* and *B. subtilis* were fixed with 4% paraformaldehyde and 1% glutaraldehyde at 4°C overnight. The fixed samples were washed with phosphate-buffered saline (PBS) and postfixed with 1% osmium tetroxide for 1 h at room temperature. After washing with PBS, the specimens were dehydrated in a graded ethanol series at room temperature. The samples were treated with propylene oxide and infiltrated with a propylene oxide-Epon (Epon 812 resin; TAAB Laboratories, Aldermaston, UK) solution [propylene oxide-Epon resin, 1:1 (v/v)] overnight. The samples were embedded in Epon resin, which was allowed to polymerize at 70°C for 72 h. Ultrathin sections were cut on an ultramicrotome (Leica Reichert Division, Vienna, Austria) and mounted on nickel grids. The sections were stained with 4% uranyl acetate and lead citrate. After staining, all sections were examined under a transmission electron microscope (model JEM1010; JEOL, Tokyo, Japan) operated at 80 kV.

### *β*-Galactosidase assay

The *β*-Galactosidase assay was performed according to the method described by Miller (38). *E. coli* cells transformed with the pGEM-T Easy plasmid were precultured in an LB medium containing 100 μg/mL ampicillin at 37°C for 14 h with reciprocal shaking (130 rpm). Precultured *E. coli* cells were inoculated to the medium with or without 5 mM diaphorin, diluting the preculture at 1:1,000, and cultured as described above. After cultivation for 4 h, isopropyl *β*-D-1-thiogalactopyranoside (IPTG) was added at a final concentration of 1 mM to induce *β*-galactosidase synthesis. Three hours after the addition of IPTG, the OD_600_ of each specimen was measured. Subsequently, 10 μL of each culture were transferred to a fresh tube and mixed with 90 μL Z buffer (60 mM Na_2_HPO_4_, 40 mM NaH_2_PO_4_, 10 mM KCl, 1 mM MgSO_4_, and 50 mM *β*-mercaptoethanol), 10 μL chloroform, and 5 μL of 0.1% sodium dodecyl sulfate solution. The tubes were vortexed and left for 1 min at room temperature to permeabilize cells. Subsequently, 20 μL of 4 mg/mL *o*-nitrophenyl-*β*-D-galactopyranoside was added to each tube. When a yellow color due to *o*-nitrophenyl developed, the reaction was stopped by adding 30 μL of 1 M Na_2_CO_3_. After centrifugation at 3,000 × *g* for 1 min, the aqueous phase was removed and used for the OD_420_ and OD_550_ measurements. The *β*-galactosidase activity was calculated using the following equations:

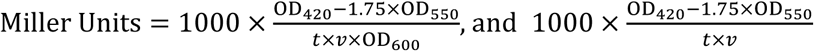

where: t = time of the enzymatic reaction (min) and v = volume of culture used in the assay (mL).

### Statistical analysis

All statistical analyses were performed using R version 4.1.3 (39). Values of bacterial cell sizes were converted into logarithms. The normal distribution of the data was assessed using the Kolmogorov-Smirnov test (40) and the Shapiro-Wilk test (41). When the normal distribution was not rejected, data from two groups were compared using Welch’s *t*-test (42). When the normal distribution was rejected, data from two groups were compared using the Brunner-Munzel test, a nonparametric method that does not assume homoscedasticity (43). For multiple comparisons, the homogeneity of variances was assessed with the Bartlett test (44). When normal distribution and homogeneous variance of data were not rejected, multiple comparisons were performed using one- or two-way analysis of variance (ANOVA), followed by Dunnett’s test (45) or Tukey’s test (46). When these null hypotheses were rejected, multiple comparisons were performed using the Kruskal-Wallis test (47), followed by the Steel test (48) or the Steel-Dwass test (49).

## Results

### Diaphorin promoted the growth of *E. coli*

To assess the effects of diaphorin on *E. coli, E. coli* strain JM109 cells were cultured in an LB medium with 100 μg/mL ampicillin supplemented with 0, 5, 50, or 500 μM or 5 mM diaphorin (Fig. 1A). Four temporally independent experiments were performed, each consisting of three independent cultures in three independent tubes per treatment, giving 12 independent cultures (n = 12) per treatment. After 6 h throughout the whole incubation time, the ΔOD_600_ of *E. coli* cultivated in a medium containing 5 mM diaphorin was significantly higher than that of *E. coli* cultured in a medium without diaphorin (*p* < 0.05, Dunnett’s test; Fig. 1A). The ratio of ΔOD_600_ of the 5 mM diaphorin group to the ΔOD_600_ of the control group reached the maximum of 1.40 at 7 h, corresponding to the logarithmic growth phase. The medium containing 5 mM diaphorin but without inoculation of *E. coli*, showed no increase in OD_600_, indicating that diaphorin does not directly affect OD_600_ in the culture medium. ΔOD_600_ of *E. coli* cultured in a medium containing 5, 50, and 500 μM of diaphorin showed no significant difference from that of *E. coli* cultured in a medium without diaphorin (*p* > 0.05, Dunnett’s test; Fig. 1A). High-throughput amplicon sequencing of the 16S rRNA gene showed that 99.992% (277,249 reads of the 277,271 total reads) and 99.995% (228,262 reads of the 228,274 total reads) of the reads derived from cultures treated with 0 and 5 mM diaphorin, respectively, corresponded to *E. coli* sequences, indicating that contamination is negligible (Supplementary Table S1).

**Figure 1.**
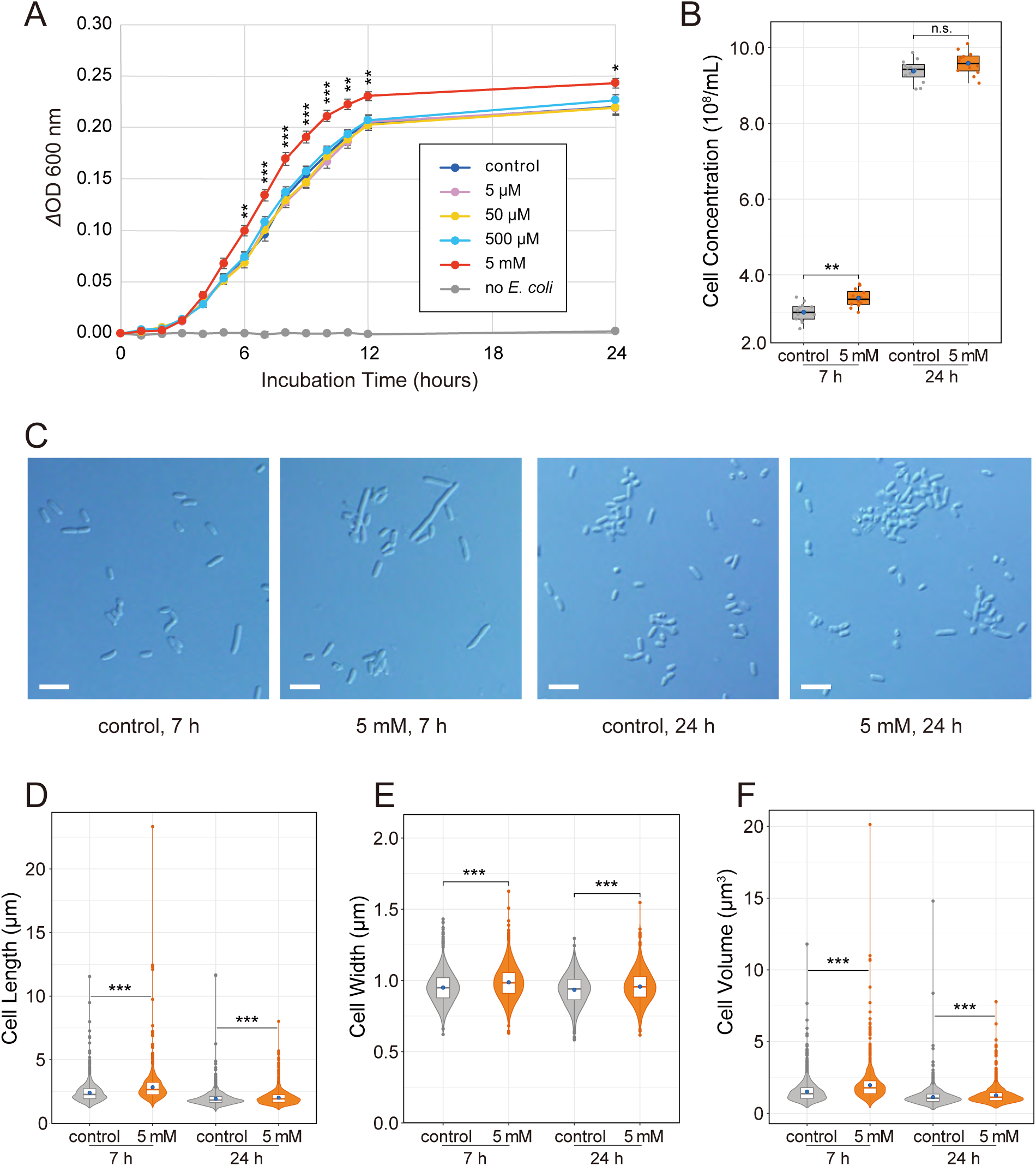
Evaluation of the biological activity of diaphorin on the growth of *E. coli*. (A) Growth dynamics of *E. coli* cultured in a medium containing 0, 5, 50, and 500 μM and 5 mM of diaphorin. The change of OD_600_ (ΔOD_600_) obtained by subtracting the value of each culture in each tube at time 0 is presented. Each data point represents the mean of 12 independent cultures (n = 12). Error bars represent standard errors (SEs). Asterisks indicate statistically significant differences (*, *p* < 0.05; **, *p* < 0.01; ***, *p* < 0.001, Dunnett’s test). To show the lack of direct effects of diaphorin on ΔOD_600_, data of a medium containing 5 mM of diaphorin but without inoculation of *E. coli* are also presented (n = 3). (B) Concentrations (numbers/mL) of *E. coli* cells cultured for 7 h (left) and 24 h (right). Jitter plots of all data points (n = 12) and box plots (gray, control; orange, 5 mM diaphorin) showing their distributions (median, quartiles, minimum, and maximum) are presented. Each data point is an average count obtained from 10 independent counting areas in a bacterial counter. Blue dots represent their means. Asterisks indicate the statistically significant difference (**, *p* < 0.01, Welch’s *t*-test) (C) DIC images of *E. coli* cultured in a medium containing 0 or 5 mM diaphorin for 7 or 24 h. Bars, 5 μm. (D) Violin plots (kernel density estimation) overlaid with box plots (median, quartiles, minimum, and maximum) and small dots (outliers) show distributions of cell lengths of *E. coli* cultured in a medium containing 0 (gray; n = 1,200) or 5 mM diaphorin (orange; n = 1,200) for 7 h (left) or 24 h (right). Blue dots represent the means. Asterisks indicate statistically significant differences (***, *p* < 0.001, Steel-Dwass test). (E) Distributions of cell widths of *E. coli* cultured in a medium containing 0 (gray; n = 1,200) or 5 mM of diaphorin (orange, n = 1,200) for 7 h (left) or 24 h (right). Symbols are the same as in (D). (F) Distributions of cell volumes of *E. coli* cultured in a medium containing 0 (gray; n = 1,200) or 5 mM of diaphorin (orange; n = 1,200) for 7 h (left) or 24 h (right). Symbols are the same as in (D).

To further examine the status of *E. coli* in these cultures, the cell concentration (numbers/mL) of cultures with and without supplementation of 5 mM diaphorin was assessed (Fig. 1B). Sampling time points were at 7 and 24 h, corresponding to the logarithmic and the stationary phases, respectively (Fig. 1A). DIC images of *E. coli* at these time points are shown in Fig. 1C. Aliquots of 12 cultures from each treatment were put into a bacterial counter, and 10 independent counting areas were used to calculate the mean concentration for each culture (Fig. 1B). At 7 h incubation, the cell concentration of cultures treated with 5 mM diaphorin was (3.38 ± 0.24) × 10^8^/mL [mean ± standard deviation (SD); n=12], which was slightly (12.3%) but significantly higher than that of control cultures at (3.01 ± 0.25) × 10^8^/mL (n = 12; *p* < 0.01, Welch’s *t*-test; Fig. 1B). At 24 h incubation, cell concentrations were not significantly different between cultures treated with 5 mM diaphorin [(9.58 ± 0.31) × 10^8^/mL (n = 12)] and control cultures [(9.38 ± 0.30) × 10^8^/mL (n = 12); *p* > 0.05, Welch’s *t*-test].

As these results showed that the increased cell concentration is not fully accountable for the observed effects of diaphorin on ΔOD_600_ of *E. coli* cultures, the morphology of *E. coli* in these cultures was subsequently assessed (Fig. 1D–F). At 7 h incubation, the length of cells treated with 5 mM diaphorin was 2.84 ± 1.18 μm (mean ± SD; n = 1,200), which was significantly larger than that of control cells at 2.41 ± 0.77 μm (n = 1,200; *p* < 0.001, Steel-Dwass test; Fig. 1D). At 24 h incubation, the length of cells treated with 5 mM diaphorin was 2.02 ± 0.56 μm (n = 1,200), which was also significantly larger than that of control cells at 1.93 ± 0.60 μm (n = 1,200; *p* < 0.001, Steel-Dwass test; Fig. 1D). Similarly, at 7 h incubation, the width of cells treated with 5 mM diaphorin was 0.99 ± 0.12 μm (n = 1,200), which was significantly larger than that of control cells at 0.95 ± 0.11 μm (n = 1,200; *p* < 0.001, Steel-Dwass test; Fig. 1E). At 24 h incubation, the width of cells treated with 5 mM diaphorin was 0.96 ± 0.11 μm (n = 1,200), which was also significantly larger than that of control cells at 0.93 ± 0.11 μm (n = 1,200; *p* < 0.001, Steel-Dwass test; Fig. 1E). Regarding cell volumes calculated from observed lengths and widths, the value of cells cultured with 5 mM diaphorin for 7 h was 1.97 ± 1.10 μm^3^ (n = 1,200), which was significantly larger (29.4%) than that of control cells at 1.52 ± 0.76 μm^3^ (n = 1,200; *p* < 0.001, Steel-Dwass test; Fig. 1F). The volume of cells cultured with 5 mM diaphorin for 24 h was 1.26 ± 0.61 μm^3^ (n = 1,200), which was slightly (9.9%) but significantly larger than that of control cells at 1.14 ± 0.63 μm^3^ (n = 1,200; *p* < 0.001, Steel-Dwass test; Fig. 1F). These results demonstrated that diaphorin, at physiological concentrations in *D. citri*, increases the concentration and cell size of *E. coli*, suggesting that diaphorin promotes the growth of *E. coli*.

### Diaphorin activated the metabolism of *E. coli*

To gain some insights into the metabolic activity of *E. coli*, the *β*-galactosidase assay was performed using *E. coli* treated with and without 5 mM diaphorin (Fig. 2; Supplementary Table S2 for values of each parameter). At the logarithmic growth phase, *E. coli* cells were incubated with 1 mM of IPTG for 3 h to induce the expression of *β*-galactosidase. The Miller unit (ΔOD_420_/min*mL*OD_600_) of *E. coli* treated with 5 mM diaphorin was 524 ± 52 (mean ± SD; n = 20), which was slightly (6.9%) but significantly larger than that of control at 490 ± 54 (n = 20; *p* < 0.05, Welch’s *t*-test; Fig. 2A). This result suggested that diaphorin activates the metabolic activity of *E. coli*. The Miller unit is based on a formula including division by OD_600_, intending to calibrate the enzymatic activity with cell density or biomass of the sample (38). When this calibration was omitted to show the enzymatic activity per culture volume, the *β*-galactosidase activity (ΔOD_420_/min*mL) of *E. coli* treated with 5 mM diaphorin was calculated to be 80.4 ± 16.9 (n = 20), which was significantly and remarkably (52.0%) larger than that of control *E. coli* at 52.9 ± 7.8 (n = 20; *p* < 0.001, Welch’s *t*-test; Fig. 2B). This result suggested that diaphorin notably activates the metabolic activity of *E. coli* per culture volume.

**Figure 2.**
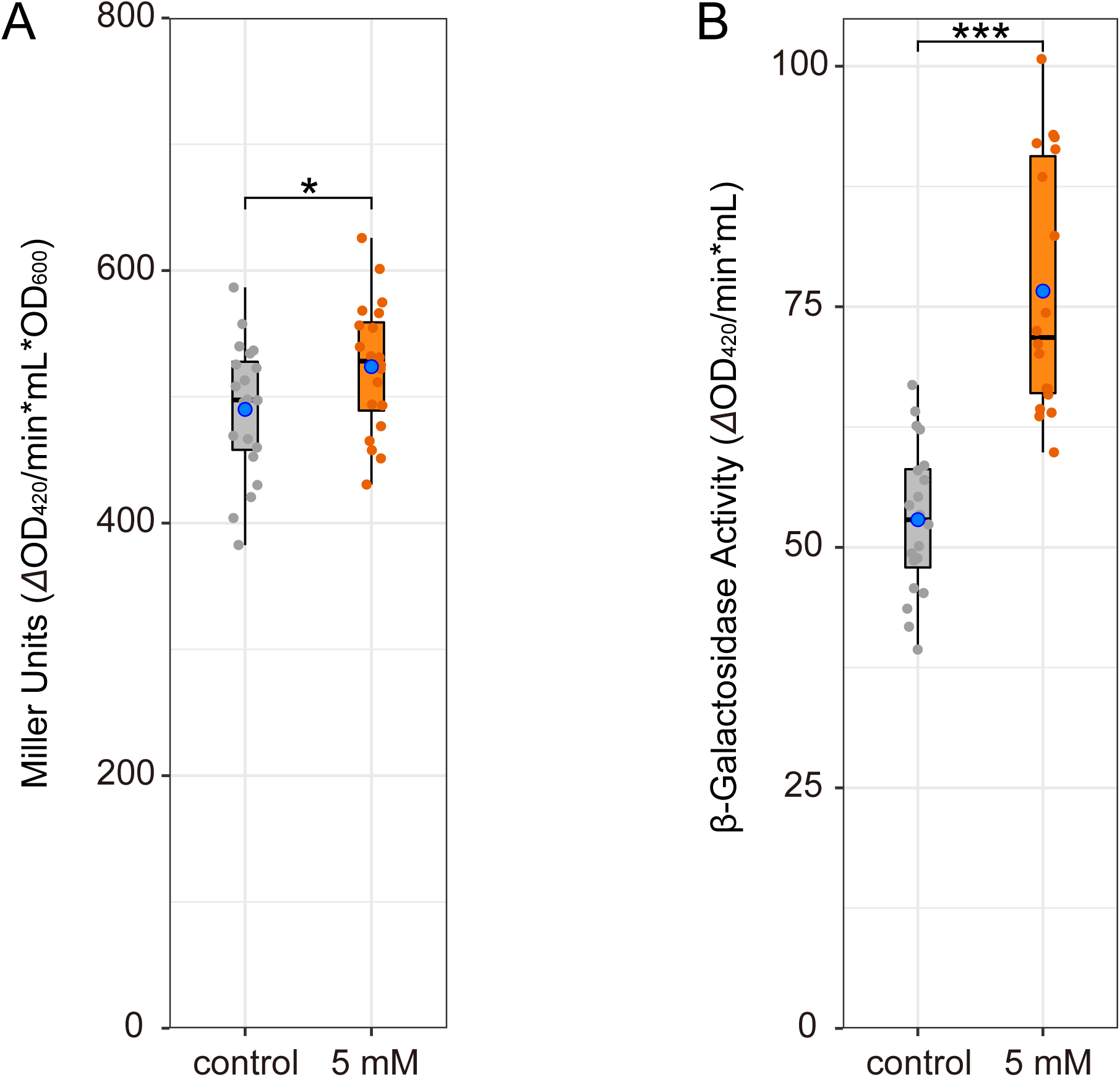
*β*-Galactosidase activity in *E. coli* cultures treated with and without diaphorin. Jitter plots of all data points (n = 20) and box plots (gray, control; orange, 5 mM diaphorin) showing their distributions (median, quartiles, minimum, and maximum) are presented. Blue dots represent the means. (A) *β*-Galactosidase activity in the form of Miller unit:

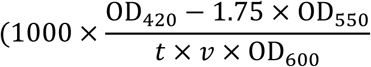

where t = time of the enzymatic reaction (min) and v = volume of culture used in the assay (mL), showing the activity relative to the cell biomass. The asterisk indicates the statistically significant difference (*, *p* < 0.05, Welch’s *t*-test). (B) *β*-Galactosidase activity without calibration with OD_600_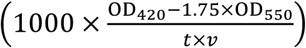, showing the activity relative to the culture volume. Asterisks indicate the statistically significant difference (***, *p* < 0.001, Welch’s *t*-test).

### Electron microscopy showed the normality of *E. coli* treated with diaphorin

To assess the ultrastructure of *E. coli* treated with diaphorin, transmission electron microscopy (TEM) was performed using *E. coli* cultured for 7 h in a medium containing 0 (Fig. 3A–D) or 5 mM of diaphorin (Fig. 3E–H). Results showed no conspicuous difference in the ultrastructure between control and diaphorin-treated *E. coli*, suggesting that *E. coli* treated with diaphorin is normal.

**Figure 3.**
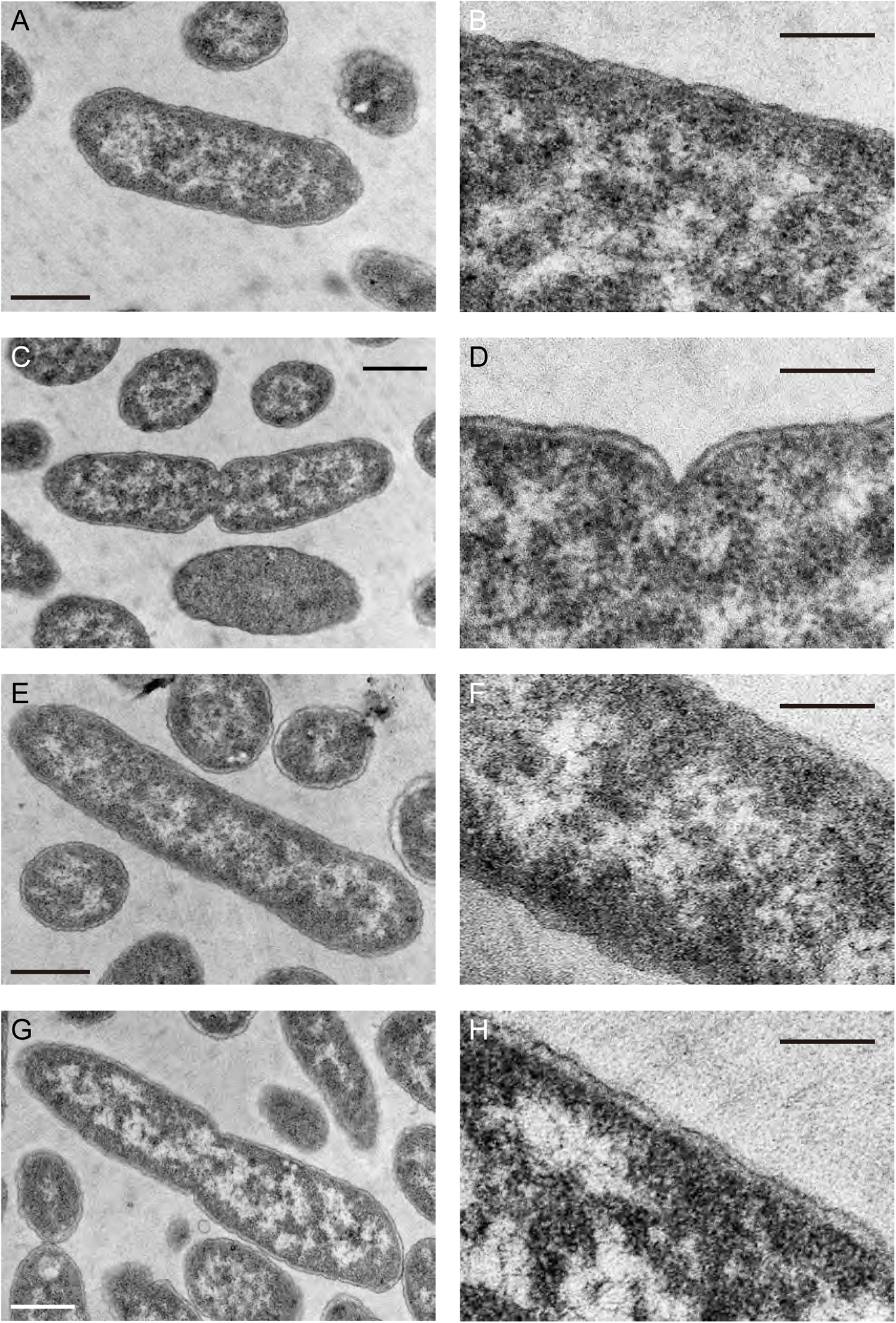
TEM of *E. coli* cultured for 7 h in a medium containing 0 (A–D) or 5 mM diaphorin (E–H). B, D, F, and H (bars, 200 nm) are magnified images of A, C, E, and G (bars, 500 nm), respectively. No conspicuous difference was observed in the ultrastructure between control and diaphorin-treated *E. coli*.

### Diaphorin inhibited the growth of *B. subtilis*

To assess the effects of diaphorin on *B. subtilis, B. subtilis* strain ISW1214 cells were cultured in an L-broth medium containing 20 μg/mL tetracycline supplemented with 0, 5, 50, or 500 μM or 5 mM diaphorin (Fig. 4A). Four temporally independent experiments were performed, each consisting of three independent cultures in three independent tubes per treatment, giving 12 independent cultures (n = 12) per treatment. After 3 h throughout the whole incubation time, the ΔOD_600_ of *B. subtilis* cultured in a medium containing 5 mM diaphorin was significantly lower than that of *B. subtilis* cultured in a medium without diaphorin (*p* < 0.001, Dunnett’s test). The ΔOD_600_ of *B. subtilis* treated with 500 μM diaphorin was also significantly lower than that of control *B. subtilis* after 5 to 24 h incubation (*p* < 0.001, Dunnett’s test). The ΔOD_600_ of *B. subtilis* cultured in a medium containing 5 and 50 μM diaphorin showed no significant difference from that of *B. subtilis* cultured in a medium without diaphorin (*p* > 0.05, Dunnett’s test). Two-way ANOVA revealed significant dosage effects of diaphorin (*F*_4, 770_ = 423.3, *p* < 0.001). The results of Tukey’s multiple comparison test are summarized in Supplementary Table S3. The medium containing 5 mM diaphorin but without inoculation of *B. subtilis* showed no increase in OD_600_ (Fig. 4A), indicating that diaphorin does not directly affect OD_600_ in the culture medium. High-throughput amplicon sequencing of the 16S rRNA gene showed that 100% of the reads (258,404 and 218,754 reads from control and 5 mM diaphorin-treated cultures, respectively) corresponded to *B. subtilis* sequences, indicating that there was essentially no contamination (Supplementary Table S4). The growth dynamics shown in this study demonstrated that diaphorin, at physiological concentrations in *D. citri*, inhibits the growth of *B. subtilis*, contrasting the case of *E. coli*.

**Figure 4.**
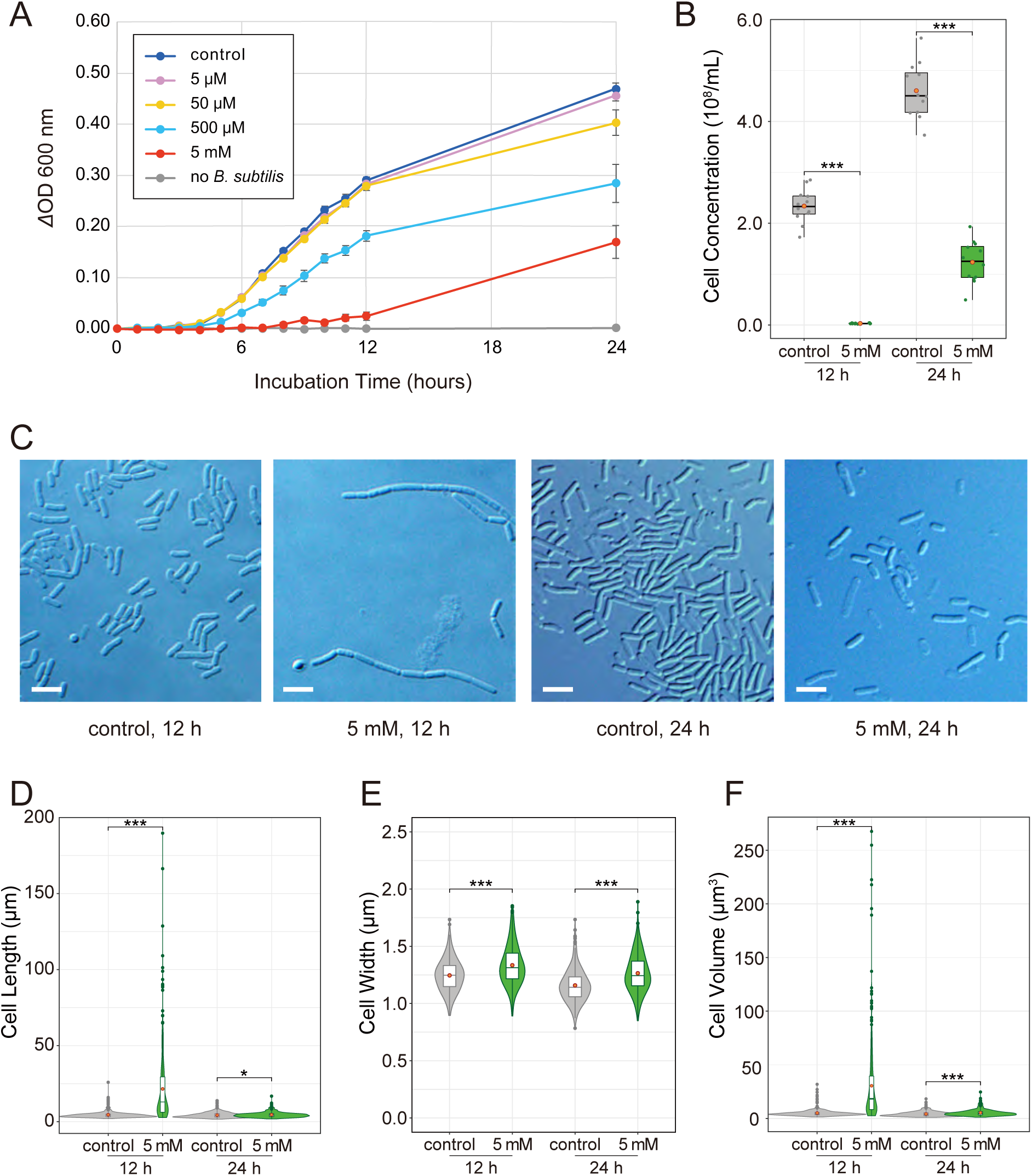
Evaluation of the biological activity of diaphorin on the growth of *B. subtilis*. (A) Growth dynamics of *B. subtilis* cultured in a medium containing 0, 5, 50, and 500 μM and 5 mM of diaphorin. The change of OD_600_ (ΔOD_600_) obtained by subtracting the value of each culture in each tube at time 0 is presented. Each data point represents the mean of 12 cultures (n = 12). Error bars represent SEs. To show the lack of direct effects of diaphorin on ΔOD_600_, data of a medium containing 5 mM of diaphorin but without inoculation of *B. subtilis* are also presented (n = 3). (B) Concentrations (numbers/mL) of *B. subtilis* cells cultured for 12 h (left) and 24 h (right). Jitter plots of all data points (n = 12) and box plots (gray, control; green, 5 mM diaphorin) showing their distributions (median, quartiles, minimum, maximum) are presented. Each data point is an average count obtained from 10 independent counting areas in a bacterial counter. Orange dots represent their means. Asterisks indicate statistically significant differences (***, *p* < 0.001, Welch’s *t*-test) (C) DIC images of *B. subtilis* cultured in a medium containing 0 or 5 mM diaphorin for 12 or 24 h. Bars, 5 μm. (D) Violin plots (kernel density estimation) overlaid with box plots (median, quartiles, minimum, maximum) and small dots (outliers) show distributions of cell length of *B. subtilis* cultured in a medium containing 0 (gray; n = 400) or 5 mM of diaphorin (green; n = 400) for 12 h (left) or 24 h (right). Orange dots represent the means. Asterisks indicate statistically significant differences (*, *p* < 0.05; ***, *p* < 0.001, Steel-Dwass test) (E) Distributions of cell widths of *B. subtilis* cultured in a medium containing 0 (gray; n = 400) or 5 mM diaphorin (green; n = 400) for 12 h (left) or 24 h (right). Symbols are the same as in (D). (F) Distributions of cell volumes of *B. subtilis* cultured in a medium containing 0 (gray; n = 400) or 5 mM diaphorin (green; n = 400) for 12 h (left) or 24 h (right). Symbols are the same as in (D).

To further examine the status of *B. subtilis* in these cultures, the cell concentration (numbers/mL) of cultures with and without supplementation of 5 mM diaphorin was assessed (Fig. 4B). Sampling time points were at 12 and 24 h. DIC images of *B. subtilis* at these time points are shown in Fig. 4C. Aliquots of 12 cultures from each treatment were put into a bacterial counter, and 10 independent counting areas were used to calculate the mean concentration for each culture. At 12 h incubation, the cell concentration of cultures treated with 5 mM diaphorin was (0.03 ± 0.00) × 10^8^/mL (mean ± SD; n = 12), which was significantly lower than that of control culture at (2.34 ± 0.33) × 10^8^/mL (n = 12; *p* < 0.001, Welch’s *t*-test; Fig. 4B). At 24 h incubation, the cell concentration of cultures treated with 5 mM diaphorin was (1.24 ± 0.41) × 10^8^/mL (n = 12), which was also significantly lower than that of control cultures at (4.60 ± 0.54) × 10^8^/mL (n = 12; *p* < 0.001, Welch’s *t*-test; Fig. 4B).

Subsequently, the morphology of *B. subtilis* cells in these cultures was assessed (Fig. 4D–F). At 12 h incubation, the length of cells treated with 5 mM diaphorin was 21.54 ± 22.82 μm (mean ± SD; n = 400), which was significantly larger than that of control cells at 4.56 ± 2.27 μm (n = 400; *p* < 0.001, Steel-Dwass test; Fig. 4D). In contrast, the length of the Hoechst-stained nucleoid area of *B. subtilis* cultured with 5 mM diaphorin for 12 h was 1.99 ± 0.59 μm (n = 400), which was significantly smaller than that of control cells at 2.51 ± 0.92 μm (n = 400; *p* < 0.001, Brunner-Munzel test; Supplementary Fig. S1), suggesting that diaphorin inhibits not only the growth but also the cleavage of *B. subtilis* cells. The length of cells cultured with 5 mM diaphorin for 24 h was 4.58 ± 1.82 μm (n = 400), which was also significantly larger than that of control cells at 4.30 ± 1.77 μm (n = 400; *p* < 0.05, Steel-Dwass test; Fig. 4D). Whereas the length of control cells was not significantly different between time points 12 and 24 h (*p* > 0.05, Steel-Dwass test), the length of cells treated with 5 mM diaphorin was significantly reduced at 24 h (*p* < 0.001, Steel-Dwass test). Regarding cell width, the value of *B. subtilis* cultured with 5 mM diaphorin for 12 h was 1.33 ± 0.17 μm (n = 400), which was again significantly larger than that of control cells at 1.24 ± 0.14 μm (n = 400; *p* < 0.001, Steel-Dwass test; Fig. 4E). At 24 h incubation, the width of cells treated with 5 mM diaphorin was 1.26 ± 0.16 μm (n = 400), which was also significantly larger than that of control cells at 1.16 ± 0.14 μm (n = 400) (*p* < 0.001, Steel-Dwass test). Diaphorin-treated and control cells significantly reduced the cell width from 12 to 24 h (*p* < 0.001, Steel-Dwass test; Fig. 4E). As for the cell volume, *B. subtilis* treated with 5 mM diaphorin for 12 h was 30.50 ± 35.51 μm^3^ (n = 400), which was significantly larger than that of control cells at 5.15 ± 3.29 μm^3^ (n = 400; *p* < 0.001, Steel-Dwass test; Fig. 4F). At 24 h incubation, the volume of cells treated with 5 mM diaphorin was 5.40 ± 3.02 μm^3^ (n = 400), which was also significantly larger than that of control cells at 4.26 ± 2.38 μm^3^ (n = 400; *p* < 0.001, Steel-Dwass test). Diaphorin-treated and control cells significantly reduced the cell volume from 12 to 24 h (*p* < 0.001, Steel-Dwass test) (Fig. 4F). These results demonstrated that diaphorin inhibits the overall growth and division of *B. subtilis* cells. The long chain of *B. subtilis* observed in this study is reminiscent of the chained cell forms in the biofilm induced by stressors, including antibiotics (50, 51). However, *B. subtilis* failed to form biofilm at 24 h incubation in this study, which may have reflected the damage to *B. subtilis* caused by diaphorin (see below). Further studies are required to understand why the chained form was temporally constructed and subsequently resolved.

### Electron microscopy showed *B. subtilis* damaged by diaphorin

To assess the ultrastructure of *B. subtilis* treated with diaphorin, TEM was performed using *B. subtilis* cultured for 12 h in a medium with and without 5 mM of diaphorin (Fig. 5). Whereas the cell envelope of control *B. subtilis* was smooth (Fig. 5A–D), the surface of cell envelopes of *B. subtilis* treated with 5 mM diaphorin was invariably rough and appeared severely damaged (Fig. 5E–J), suggesting the harmful effects of diaphorin on the cell envelope of *B. subtilis*. Additionally, “mesosome”-like structures were frequently observed in *B. subtilis* cells treated with diaphorin (Fig. 5E and F). In some extreme cases, cells were filled with membranous structures similar to mesosomes (Fig. 5I and J). These membranous structures were not conspicuous in control *B. subtilis* (Fig. 5A–D). Mesosomes, which are intracytoplasmic membrane inclusions or invagination of the plasma membrane, are recognized to be structural artifacts induced by chemical fixatives used to prepare electron microscopic specimens (52). However, such structures are often preferentially observed in bacteria treated with antimicrobial agents, including antibiotics and antimicrobial peptides (53–55), indicative of alterations in the cytoplasmic membranes caused by these agents. In this study, high levels of extent and frequency of “mesosome”-like membranous structures were observed only in diaphorin-treated *B. subtilis*, implying that these ultrastructures reflect the actual effects of diaphorin on *B. subtilis*.

**Figure 5.**
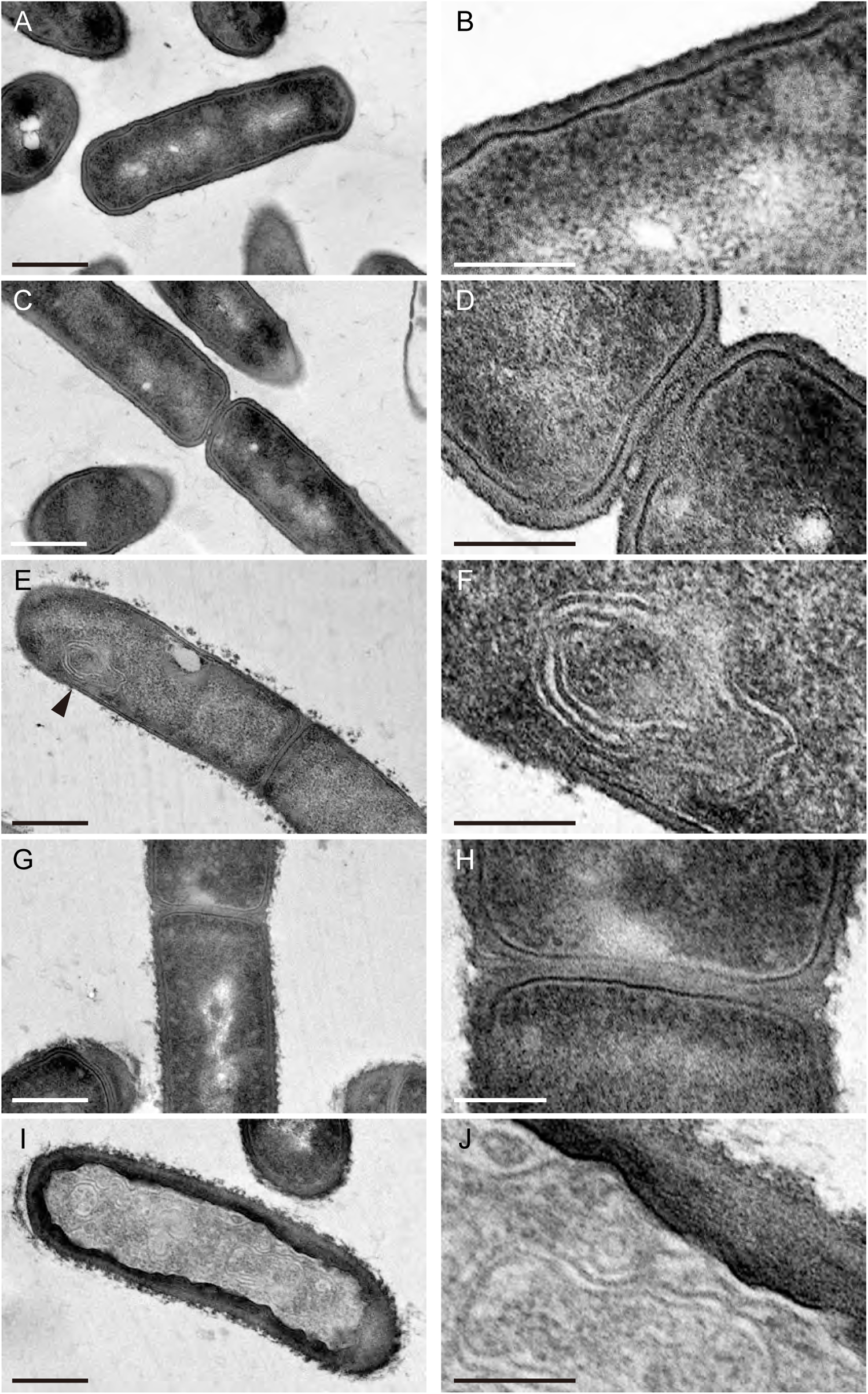
TEM of *B. subtilis* cultured for 12 h in a medium containing 0 (A–D) or 5 mM of diaphorin (E–J). B, D, F, H, and J (bars, 200 nm) are magnified images of A, C, E, G, and I (bars, 500 nm), respectively. Whereas the cell envelope of control *B. subtilis* was smooth (A–D), the surface of cell envelopes of *B. subtilis* treated with diaphorin was invariably rough and appeared disrupted (E–J), suggesting the harmful effects of diaphorin on the *B. subtilis* cell envelope. “Mesosome”-like structures were observed in *B. subtilis* cells treated with diaphorin (E; arrowhead, F). In some cases, cells were filled with cytoplasmic membranous structures similar to mesosomes (I and J).

## Discussion

The present study revealed that the physiological concentration of diaphorin, a polyketide synthesized by an obligate symbiont of psyllids, inhibits the growth of *B. subtilis* (Gram-positive bacteria) but promotes the growth of *E. coli* (Gram-negative bacteria). As exemplified by some antibiotics, certain secondary metabolites have inhibitory effects only on Gram-positive bacteria that lack the outer membrane, an effective barrier that protects Gram-negative bacteria from toxic compounds (2, 4). However, it is unique that a single molecule clearly exhibits opposite effects on distinct bacterial lineages. Particularly, the observed positive effects of diaphorin on *E. coli* attract the authors’ interest. As mentioned above, *D. citri* has two bacterial symbionts, *Ca*. Carsonella ruddii (Gammaproteobacteria: Oceanospirillales), and *Ca*. Profftella armatura (Gammaproteobacteria: Burkholderiales) (21, 22). Additionally, many populations of *D. citri* are infected with *Wolbachia* (Alphaproteobacteria: Rickettsiales), a potential manipulator of host reproduction (12, 31, 56, 57). Moreover, some *D. citri* populations are infected with *Ca*. Liberibacter spp. (Alphaproteobacteria: Rhizobiales), the causative agents of the citrus greening disease, HLB (9–12, 57). Although *Liberibacter* was shown to reduce the nymphal development rate and adult survival, it was also demonstrated to increase the fecundity, female attractiveness to males, and propensity for dispersal of *D. citri* (58, 59). Thus, this bacterial lineage can be beneficial for psyllid vectors in some ecological contexts. As with cases in other hemipteran insects (60–71), recent studies are revealing that not only interactions between host psyllids and symbiotic microbes, including those associated with the bacteriome, facultative symbionts, and plant pathogens (19–23, 29), but also interactions among such bacterial populations are important for psyllid biology and host plant pathology (11, 12, 22, 31, 35, 72). Interestingly, all the above-mentioned symbionts in *D. citri*, namely, *Carsonella, Profftella, Wolbachia*, and *Liberibacter*, belong to the phylum Proteobacteria and are closely related to *E. coli*, on which diaphorin exhibited positive effects. The bacteriome-associated obligate symbionts, *Carsonella* and *Profftella*, are especially close relatives of *E. coli*, all belonging to the class Gammaproteobacteria. Thus, it would not be farfetched to assume that diaphorin may potentially have positive effects also on these bacterial symbionts, protecting the holobiont from other intruding bacterial lineages on the other hand. Moreover, in the present study, the *β*-galactosidase assay suggested that diaphorin activates the metabolic activity of *E. coli*. As *E. coli* is utilized for producing various industrially important materials, including pharmaceutical drugs, amino acids, enzymes, and biofuels (73–76), these observed effects of diaphorin may be exploited to promote the efficiency of industrial material production by *E*.*coli*. To pursue this possibility, further studies are warranted to understand the detailed target microbial spectrum and elaborate on the mechanisms for the exhibited biological activities of diaphorin.

Also, in pest management, the target spectrum of diaphorin potentially affects the effectiveness of the biological control of *D. citri* using entomopathogenic bacteria. A notable report on *D. citri* exposed to bacteria (77) which showed that Gram-negative bacteria, including *E. coli*, significantly increased the mortality of *D. citri*, but Gram-positive bacteria, including *B. subtilis*, did not. During the experiment, *E. coli* titers increased rapidly after exposure and remained high until the death of *D. citri* (77), which appeared consistent with the fact that *D. citri* lacks genes for the Imd pathway (78), an immune pathway targeting Gram-negative bacteria with diaminopimelic acid (DAP)-type peptidoglycan (79). In contrast, *D. citri* has a nearly complete Toll immune pathway targeting Gram-positive bacteria with lysine-type peptidoglycan (78). However, *B. subtilis*, the model Gram-positive bacterium, has DAP-type peptidoglycan in its cell wall like Gram-negative bacteria and is exclusively recognized by the Imd pathway (80). Thus, it was an enigma why exposure to *B. subtilis* caused no damage to *D. citri*, which lacks the Imd pathway and most genes for antimicrobial peptides (77). The inhibitory effects of diaphorin on *B. subtilis*, demonstrated in the present study, appear to provide the answer to this enigma.

## Conclusion

The preset study revealed that diaphorin (1) inhibits the growth of *B. subtilis* and (2) promotes the growth of *E. coli*. This finding provides insights into the potential role of diaphorin in facilitating symbiotic associations, manipulating bacterial populations within *D. citri*. This can also be exploited to promote the effectiveness of industrial material production by microorganisms. Further studies are required to reveal the biological activities of diaphorin on more diverse bacterial lineages and the molecular mechanisms for exerting observed activities.

## Acknowledgments

This work was supported by the Japan Society for the Promotion of Science (https://www.jsps.go.jp) KAKENHI (grant numbers 26292174 and 20H02998), the NIBB Collaborative Research Program for Integrative Imaging (21-417), and research grants from Tatematsu Foundation and Nagase Science and Technology Foundation to AN. The funders had no role in study design, data collection and analysis, decision to publish, or preparation of the manuscript.

## Figure legends

**Figure S1**. Chained *B. subtilis* cells caused by diaphorin treatment. (A) DIC and fluorescence images of Hoechst-stained *B. subtilis* cells after cultivation for 12 h in a medium containing 0 or 5 mM of diaphorin. Bars, 5 μm. (B) Lengths of Hoechst-stained nucleoid areas in *B. subtilis* cells cultured in a medium containing 0 (gray; n = 400) or 5 mM diaphorin (green; n = 400) for 12 h. Violin plots showing kernel density estimation are overlaid with box plots showing median, quartiles, minimum, maximum, and outliers. Orange dots represent the means.

